# Sequential Operations Revealed by Serendipitous Feature Selectivity in Frontal Eye Field

**DOI:** 10.1101/683144

**Authors:** Kaleb A. Lowe, Jeffrey D. Schall

## Abstract

Neurons in macaque frontal eye field contribute to spatial but typically not feature selection during visual search. Using an innovative visual search task, we report a serendipitous discovery that some frontal eye field neurons can develop rapid selectivity for stimulus orientation that is used to guide gaze during a visual search task with pro-saccade and anti-saccade responses. This feature selectivity occurs simultaneously at multiple locations for all objects sharing that feature and coincides with when neurons select the singleton of a search array. This feature selectivity also reveals the distinct, subsequent operation of selecting the endpoint of the saccade in pro-saccade as well as anti-saccade trials. These results demonstrate that target selection preceding saccade preparation is composed of multiple operations. We conjecture that singleton selection indexes the allocation of attention, which can be divided, to conspicuous items. Consequently, endpoint selection indexes the focused allocation of attention to the endpoint of the saccade. These results demonstrate that saccade target selection is not a unitary process.

**SIGNIFICANCE STATEMENT:** Frontal eye field is well known to contribute to spatial selection for attention and eye movements. We discovered that some frontal eye field neurons can acquire selectivity for stimulus orientation when it guides visual search. The chronometry of neurons with and without feature selectivity reveal distinct operations accomplishing visual search.

## INTRODUCTION

To navigate in and interact with the visual world, primates must locate and identify objects to scrutinize through gaze. To understand how this localization, identification and gaze shifting is performed, we use visual search tasks in which targets for gaze shifts are presented with distracting stimuli. Target stimuli can be distinguished from distractors by some feature or set of features (Wolfe & Utochkin, 2019). Targets are sought through an interplay of localization, identification, and saccade preparation manifest as covert and overt orienting.

The frontal eye field (FEF), in prefrontal cortex, is known to support attention and eye movements and the performance of visual search (see Bisley & Mirpour, 2019; Schall, 2015 for review). Neurons in FEF respond to visual stimulation, before eye movements, or both (Bruce & Goldberg, 1985; Lowe & Schall, 2018; Schall, 1991). FEF has been conceptualized as a salience or priority map (Bisley, 2011; Fernandes et al., 2014; Thompson & Bichot, 2005), meaning that its responses are related to whether a stimulus is important for attention or gaze shifts regardless of what features make it important (Mohler et al., 1973; Monosov et al., 2010; Ogawa & Komatsu, 2006; Ramkumar et al., 2016; Schall et al., 1995; Zhou & Desimone, 2011). However, FEF is also an ocular motor center (Schall, 2015). Therefore, experimental manipulations are needed to dissociate selection of a stimulus as a conspicuous object, selection of a stimulus as a potential endpoint of a gaze shift, or preparation of a saccade (Matsushima & Tanaka, 2014; Murthy et al., 2001; Sato et al., 2001; Sato & Schall, 2003; Scerra et al., 2019; Thompson et al., 1996; Trageser et al., 2008; c.f. Costello et al., 2013).

Our laboratory designed a visual task to dissociate localization of a color singleton from the endpoint of a saccade reporting its location (Sato & Schall, 2003; Schall, 2004). The orientation of a color singleton cued monkeys to produce either a pro-saccade to the singleton or an anti-saccade to the distractor at the opposite location. We have improved the task by making the distractors elongated. This requires monkeys to select on color but respond on shape, resembling classic filtering tasks (Eriksen & Eriksen, 1974; Sperling, 1960; Theeuwes, 1992; Treisman & Gelade, 1980). The literature is divided on whether selecting an object and categorizing it are separate, sequential stages (Broadbent, 1971; Hoffman, 1978; Treisman, 1988; Wolfe et al., 2015) or objects are selected and categorized in a single step (Bundesen, 1990; Logan, 2002). Thus, whether covert and overt orienting processes are comprised of distinct operations or stages remains uncertain.

These differing views can be resolved through measurements of neural chronometry (Fig. 1). In the pursuit of this research aim, reward contingencies allowed one monkey to discover a strategy that prioritized the shape of the stimuli. Unexpectedly, some neurons recorded during this task exhibited rapid selectivity for stimulus shape. Here, we compare these findings to a previous report of color selectivity in FEF (Bichot et al., 1996) and characterize the neural chronometry of these FEF neurons. The results provide new evidence that selection of objects and saccade endpoints are distinct operations, both accomplished by visually responsive FEF neurons. The time course of this feature selectivity provides new evidence that visual search is accomplished through sequential operations.

**Figure 1.**
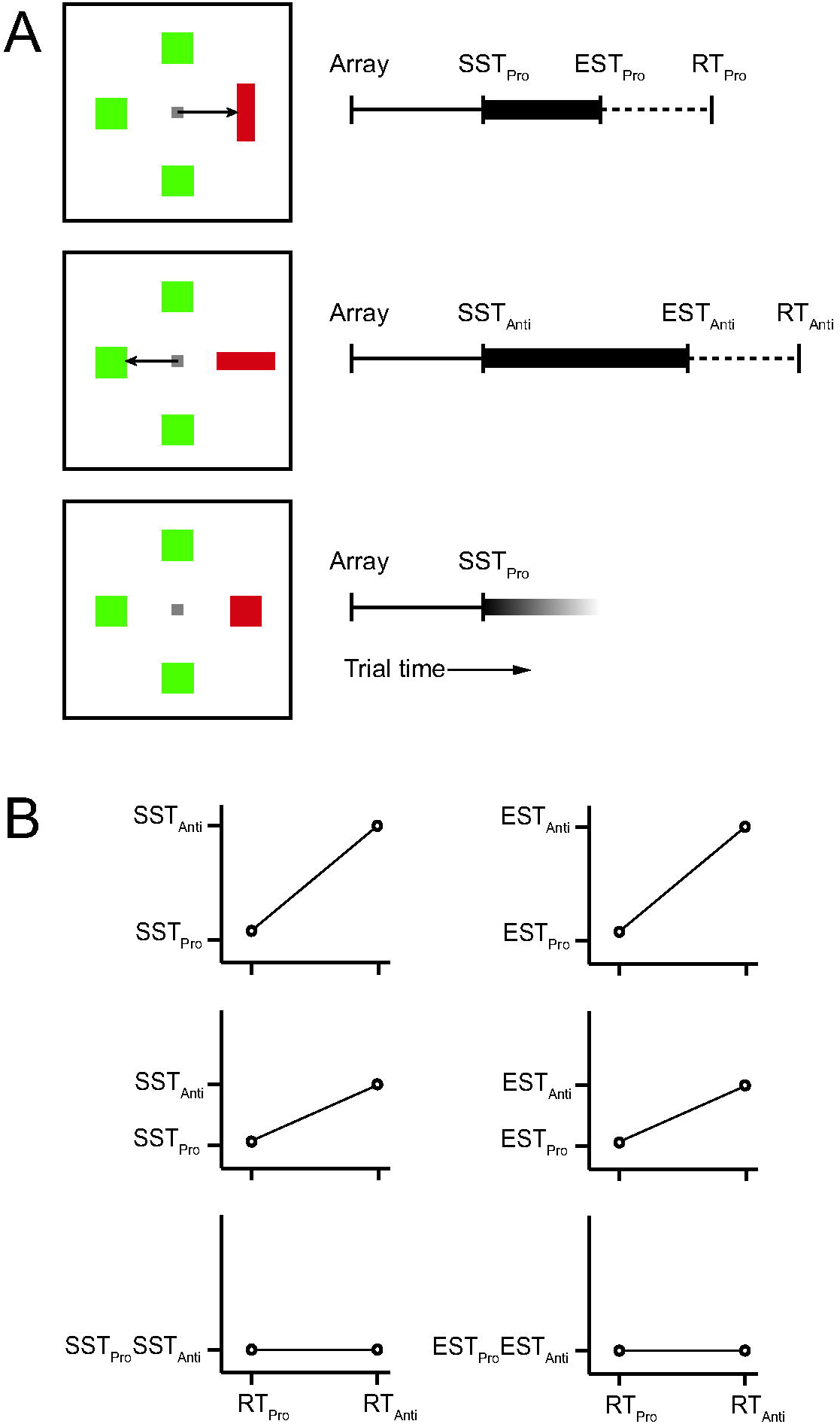
Visual search with explicit stimulus-response mapping. (A) Visual search task in which the orientation of a color singleton cues a pro-saccade (vertical), an anti-saccade (horizontal), or no saccade (square). Response times can be subdivided into three states or operations. Array presentation is followed by stimulus encoding and localization (thin line); the conclusion of this operation is indexed by singleton selection time (SST). Next, stimulus-response mapping and selection of the saccade endpoint happens if a pro- or anti-saccade will be produced (thick line); the conclusion of this operation is indexed by endpoint selection time (EST). This operation may not occur when no saccade is made (grayed thick line). Finally, saccade preparation leads to initiation of the saccade which is manifest as the measurement of RT (dotted line). (B) Response time on anti-saccade trials (RT_Anti_) is systematically longer than that on pro-saccade trials (RT_Pro_). Measurements of SST and EST provide insight into the operations contributing to the variation of RT. Theoretically, a difference between SST_Anti_ and SST_Pro_ (left) or between EST_Anti_ and EST_Pro_ (right) could explain all (top), some (middle), or none (bottom) of the variation of RT.

## METHODS

### Subjects

Data from one male macaque monkey (*M. radiata*) was compared to data previously collected from four male macaque monkeys (*M. mulatta*). All procedures were in accordance with the National Institutes of Health Guide for the Care and Use of Laboratory Animals and approved by the Vanderbilt Institutional Animal Care and Use Committee.

### Visual Search Task

All macaque monkeys performed color singleton visual search tasks. For two monkeys (A, C) the colors of singleton and distractor were constant, giving rise to strong search performance asymmetries (Bichot et al., 1996). For two monkeys (B, Q) the singleton and distractors alternated between red/green or green/red across sessions. New performance and neurophysiology data were collected from another monkey (Da) performing the visual search task with pro- and anti-saccades (Sato & Schall, 2003). The orientation of the singleton cued the pro- or anti-saccade and was presented with elongated distractors. The monkey was trained to fixate a central point whose appearance marked the beginning of the trial. After fixating this point for between 300 and 800 ms, an array of four rectangular stimuli appeared between 3° and 10° eccentricity. One of these stimuli was a color singleton (either red with green distractors or green with red distractors). The color of the singleton and distractors were randomly assigned on a trial by trial basis. All stimuli had an area of 1 square degree. Singletons could be either vertical (aspect ratio = 4.00) or horizontal (aspect ratio=0.25). Distractors could be either vertical, horizontal, or square (aspect ratio = 1.00). The aspect ratio of the color singleton indicated a response rule. If the singleton was vertical then reward was delivered for a saccade to the singleton (pro-saccade; Fig. 2A). If the singleton was horizontal then reward was delivered for a saccade to the stimulus located opposite to the singleton (anti-saccade). After making the saccade, the monkey was required to fixate the correct stimulus for 400 ms, until the fluid reward was delivered. If the monkey broke fixation or made a saccade to an incorrect location, a 2,000 ms time-out delay occurred.

**Figure 2.**
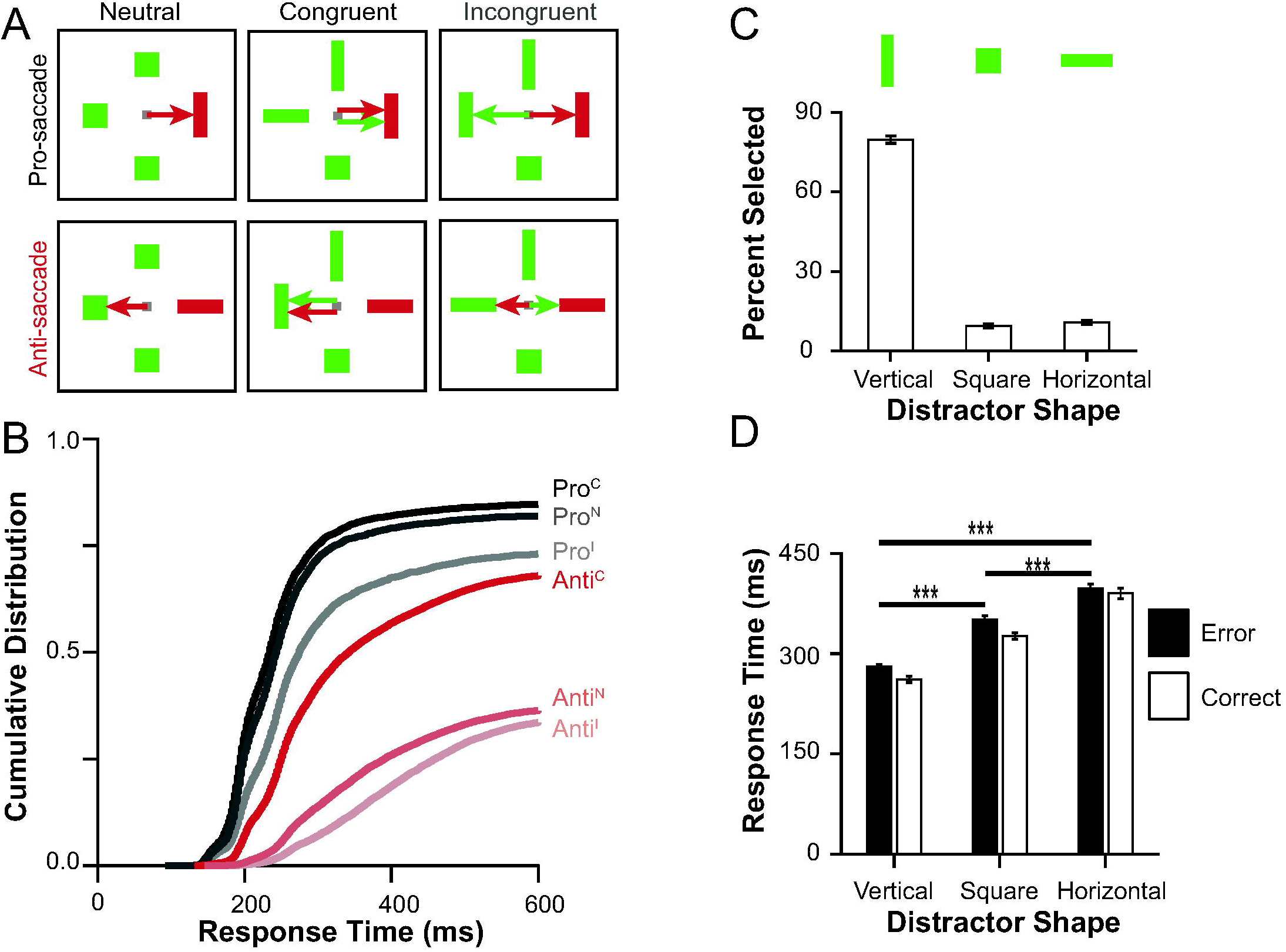
Search array configurations and task performance. (A) Visual search with pro-saccade (top) and anti-saccade (bottom) responses based on orientation of color singleton. Distractors could be square or elongated. Because shape of the singleton cues stimulus-response rule, the shape of the distractors may influence the efficiency of stimulus-response mapping via a congruency effect. We operationalized neutral trials as those in which the distractor opposite the singleton was square (left column), congruent trials as those in which the distractor opposite the singleton would cue the same saccade as the singleton (middle column), and incongruent trials as those in which the distractor opposite the singleton cued the opposite saccade (right column). The saccades cued by the singleton (distractor) are indicated as red (green) arrows. (B) Defective RT distributions for pro-saccade (black) and anti-saccade (red) trials with congruent arrays (full saturation), neutral arrays (intermediate saturation), and incongruent arrays (lowest saturation). Saccade latency was longer for anti-relative to pro-saccades, and longer of incongruent relative to neutral and congruent trials. (C) Proportions of error saccades made to each stimulus shape for trials in which at least one distractor was vertical (open). (D) RTs to each stimulus shape for error (filled) and correct (open) trials. Saccades to vertical items were shortest latency.

Correct responses were defined by the orientation of the color singleton. Hence, the orientation of the distractors can influence response selection. Consequently, particular combinations of singleton and distractor orientations can cue congruent or incongruent saccades. The distractor opposite the singleton was a correct endpoint on anti-saccade trials, so congruency was operationalized by the relationship of the shape of the singleton and the distractor at the opposite location. If the distractor was vertical, a saccade may be planned toward it. If it was horizontal a saccade may be planned toward the color singleton. If the saccade consistent with the orientation of the opposite distractor corresponded to the saccade cued by the singleton, then the stimulus array was *congruent*. If the singleton and opposite distractor cued saccades in opposite directions, then the stimulus array was *incongruent*. If the opposite stimulus was square, the stimulus array was *neutral*.

### Data acquisition and analysis

Because all details have been described previously (Cohen et al., 2009; Sato et al., 2001; Schall et al., 1995), they will not be repeated. The following approaches and definitions are particular to this analysis.

For averaging across neurons, SDFs were normalized by z-scoring across the full trial and performing a baseline subtraction. That is, the SDFs aligned on array presentation and saccade for each condition were concatenated and the standard deviation of this concatenated vector was calculated. The SDFs for that unit were then divided by that standard deviation.

Then, the mean baseline activity, the average value of the SDF in the 300 ms preceding array onset, was subtracted. This method of scaling responses reduces the skewness of the SDF across the population and generates a comparable range of activity across neurons without erroneously scaling neurons with little to no modulation (Lowe & Schall, 2018).

Selection times were calculated from the SDFs by subtracting the mean difference during the 300 ms before array onset from the difference between two conditions. Selection times were defined as the earlier of two times (1) the time the difference function exceeds 2 standard deviations of the baseline difference and continues on to exceed 6 standard deviations for at least 20 ms continuously or (2) the time the difference function exceeds 2 standard deviations of the baseline difference for at least 50 ms continuously. Visual latency was calculated in a similar fashion where the SDF itself meeting the above criteria as opposed to a difference function. Differences among selection time distributions were assessed with a nonparametric Kruskal-Wallis test for equal medians.

Each selection time measure was calculated over all RTs and in groups of trials with shortest and longest RTs based on median split. The magnitude of any difference in selection times across RT groups was compared to the difference in RT across the groups through a two-tailed t-test and associated Bayes factor.

## RESULTS

### Performance Results

We begin by introducing a nomenclature used below. Correct saccades to vertical stimuli included pro-saccade trials with congruent, neutral, or incongruent arrays (Pro^C,N,I^) and congruent anti-saccade trials (Anti^C^). We also designate saccades to square stimuli as neutral anti-saccade trials (Anti^N^) and saccades to horizontal stimuli as incongruent anti-saccade trials (Anti^I^).

RT and accuracy both exhibited an influence of response mapping and singleton-distractor congruency (Fig. 2B). As expected, mean RT ± SEM on all anti-saccade trials (311 ± 48 ms) was significantly greater than RT on all pro-saccade trials (240 ± 28 ms) (ANOVA: F(1,198) = 182.5, p < 0.001. A Bayesian analysis suggested that the data were 2.8 x 10^22^ times as likely to have been observed in a model including stimulus-response mapping as a factor as compared to a null model. Also, RT on all incongruent trials (304 ± 57 ms) was significantly greater than RT on all neutral trials (282 ± 50 ms), which was significantly greater than RT on all congruent trials (260 ± 45 ms) (ANOVA: F(2,198) = 20.9, p < 0.001). A Bayesian analysis suggested that the data were 1.7 x 10^7^ times as likely to have been observed in a model including congruency in addition to stimulus-response mapping as compared to a model with stimulus-response mapping alone. Thus, the shape of the distractors influenced the efficiency of visual search and saccade production. A Bayesian analysis suggested no evidence of an interaction; the data were 1.24 times as likely to have been observed in a model with no interaction as compared to a model with an interaction between stimulus-response mapping and congruency.

Analyzing the pattern of errors, we discovered that the monkey more commonly shifted gaze to a vertical item than to any other (Fig. 2C). Endpoint errors were significantly more common to vertical stimuli (80 ± 12% vertical, 10 ± 7 % square, 11 ± 7% horizontal; ANOVA: F(2,117) = 833.92, p < 0.001). A Bayesian analysis suggested that the data were 3.6 x 10^65^ times as likely to have been observed in a model including shape as a factor as compared to a null model. The preference for vertical stimuli was evident also in the RT (Fig. 2D). RTs were significantly shorter for saccades to vertical (271 ± 38 ms), relative to square (339 ± 49 ms) and horizontal stimuli (394 ± 67 ms) (ANOVA: F(2,234) = 110.15, p < 0.001) regardless of correct or error trial outcome (ANOVA: interaction F(2,234) = 0.58, p = 0.561). A Bayesian analysis suggested that the data were 3.8 x 10^30^ times as likely to have been observed in a model including shape as a factor as compared to a null model. There was also no evidence of an interaction, as the data were 8.3 times as likely to have been observed in a model with only shape and trial outcome as factors as compared to a model with an interaction. The more frequent and faster responses to vertical stimuli indicate that the monkey adopted a strategy of searching for vertical items as opposed to guiding gaze by the stimulus-response rule provided by the singleton. In other words, the monkey divided attention to vertical items in the array rather than focusing attention on the singleton that cued the stimulus-response rule. Serendipitously, the short-cut used by the monkey revealed new properties of feature and spatial processing supporting visual search with arbitrary stimulus-response mapping.

### Shape Selectivity in FEF

Based on previous observations during color singleton search with fixed target and distractor color assignments (Bichot et al. 1996), we tested whether the predisposition for vertical stimuli was associated with altered stimulus feature processing by FEF neurons. FEF is comprised of a diversity of neurons with visual, visuomovement, movement, and other patterns of modulation (Lowe & Schall 2018). The sample of neurons analyzed for this report consisted entirely of visually responsive neurons. This is important to understand because we will describe a pattern of modulation that is related to saccade production but is distinct from the saccade preparation accomplished by movement neurons.

Responses to the different stimulus shapes was assessed when they were irrelevant distractors, i.e., not the color singleton nor the endpoint of an anti-saccade or error saccade. Responses to vertical, square, and horizontal irrelevant distractors from two example neurons are shown in Fig. 3A. Both neurons responded more to a vertical than to any other item in the RF. The time at which this difference between responses to vertical and non-vertical stimuli was defined as *feature selection time (*FST). For neuron 1, FST occurred 136 ms after array presentation, 41 ms after the initial transient. FST for neuron 2 occurred 95 ms after array presentation, only 8 ms after the visual transient. These representative neurons exemplify two other distinctive properties. Whereas neuron 1 showed graded selectivity (vertical > square > horizontal), neuron 2 showed categorical selectivity (vertical > square = horizontal) (e.g., Ferrera et al., 2009). The average responses to vertical, square, and horizontal objects for the feature selective neurons is shown in Fig. 3B. The mean ± SEM FST was 130 ± 30 ms (mode = 134 ms; Table 1).

**Figure 3.**
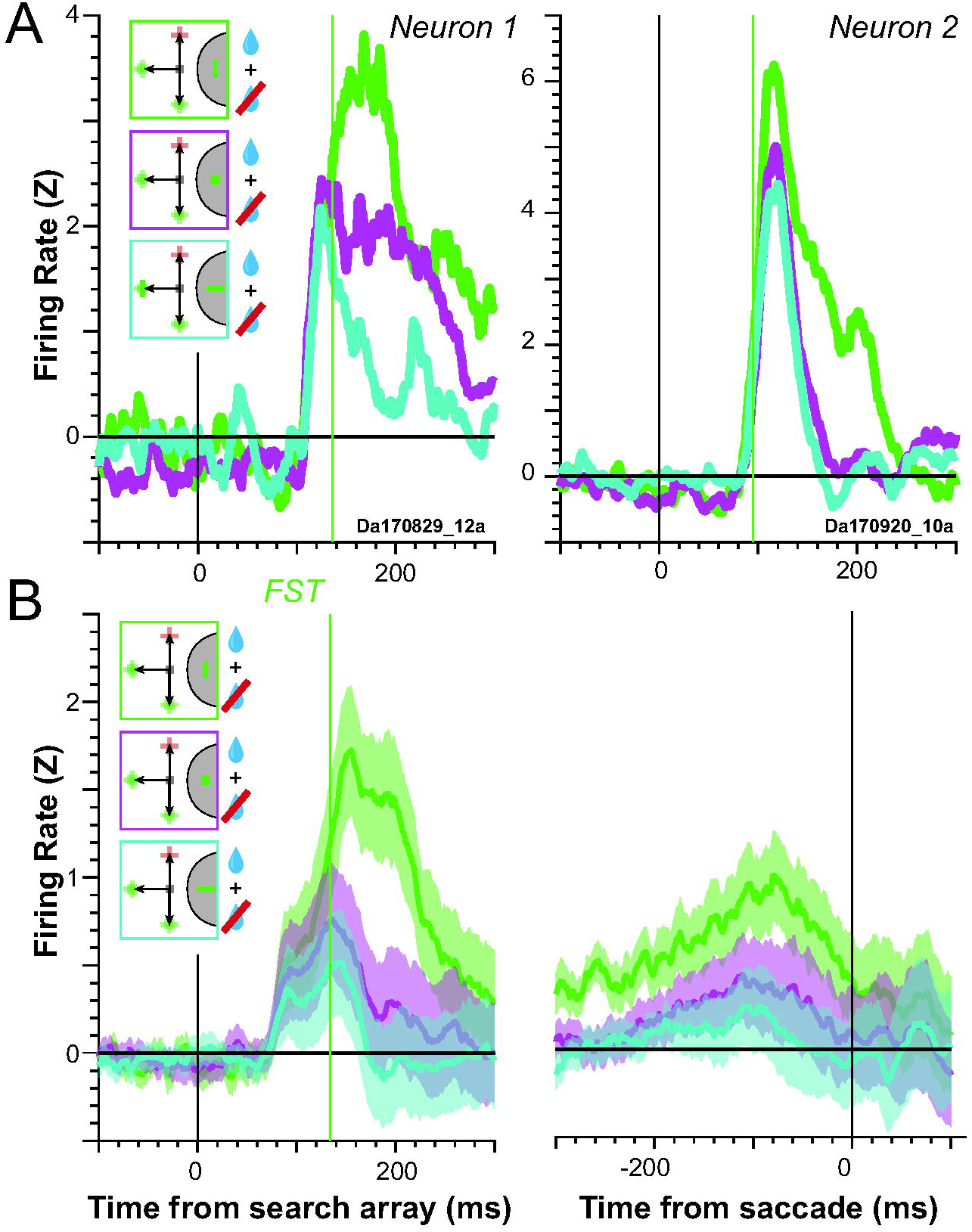
Feature selectivity in FEF. (A) Normalized firing rate for two example neurons that exhibited shape selectivity aligned on stimulus onset. Responses to vertical (green), square (magenta), and horizontal (cyan) stimuli that were irrelevant distractors across correct (blue drop) and error (crossed blue drop) pro-and anti-saccade trials. Trial types are indicated in the color-coded insets. The set of possible stimuli that can appear at a given location are superimposed. The singleton shown at 90° could have appeared at 270°; likewise, the distractors shown at 270° could have appeared at 90°. Feature selection time (FST) is indicated by the vertical green line. (B) Average normalized firing rate ± SEM for all feature selective neurons aligned on array presentation (left) and saccade initiation (right). Vertical green line plots the median FST for this population.

**Table 1.**
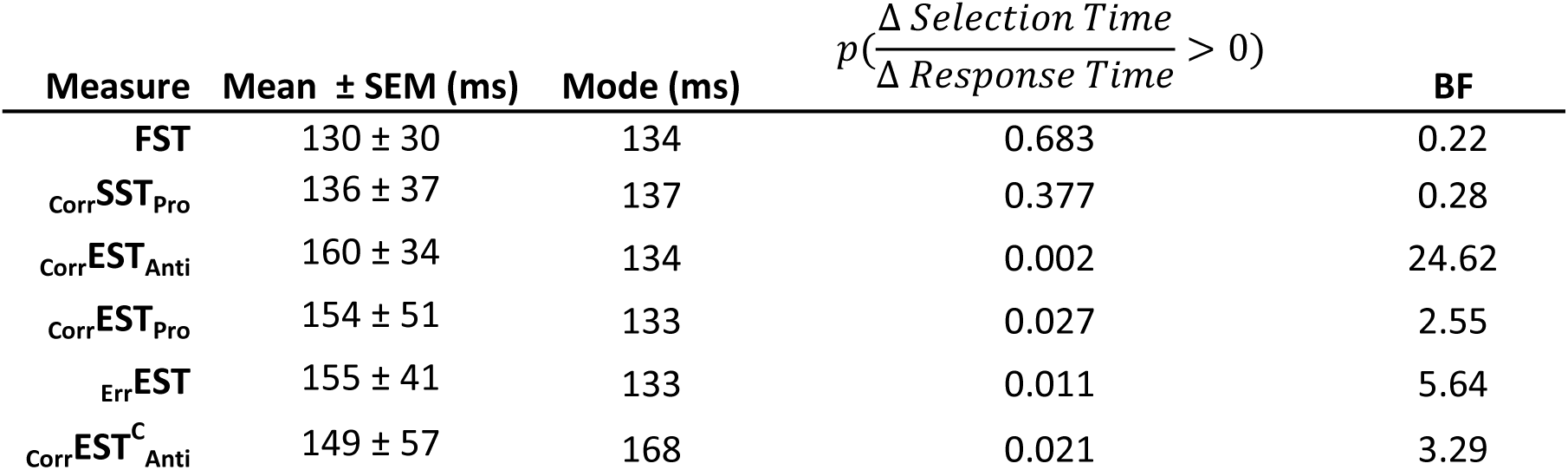
Selection time summary statististics. For each selection time, the table reports the mean value ± SEM, modal value, probability that variation in selection time over interquartile range of the response times is equal to zero (i.e., the probability that selection time is synchronized to array presentation), and the Bayes factor for whether the change in selection time is synchronized to the change in RT (BF < 0) or not synchronized to the change in RT (BF > 0).

In monkeys performing color singleton search with constant target and distractor colors, the color-selective neurons in FEF responded with latencies not less than ∼60 ms, while non-selective neurons responded with latencies as short as ∼40 ms (Bichot et al., 1996). We compared the current results to those data (Fig. 4). For each neuron, an ANOVA was performed on the SDF values during the first 25 ms (corresponding to the interval used by Bichot et al. (1996)) or 100 ms after the visual transient. Of 124 neurons sampled, 13% showed shape selectivity in the first 25 ms and 24% in the first 100 ms. As observed previously, neurons with shape selectivity were not the earliest to respond. The earliest visual response of shape selective neurons was 66 ms (median 95 ms; mode 89 ms), later than the two earliest visual responses from non-shape selective neurons 52 and 58 ms). Combined across the two studies, the results show that neither shape nor color information arrives in FEF via the fastest visual pathway and indicate that the training conditions of the present study created the same feature selective state.

**Figure 4.**
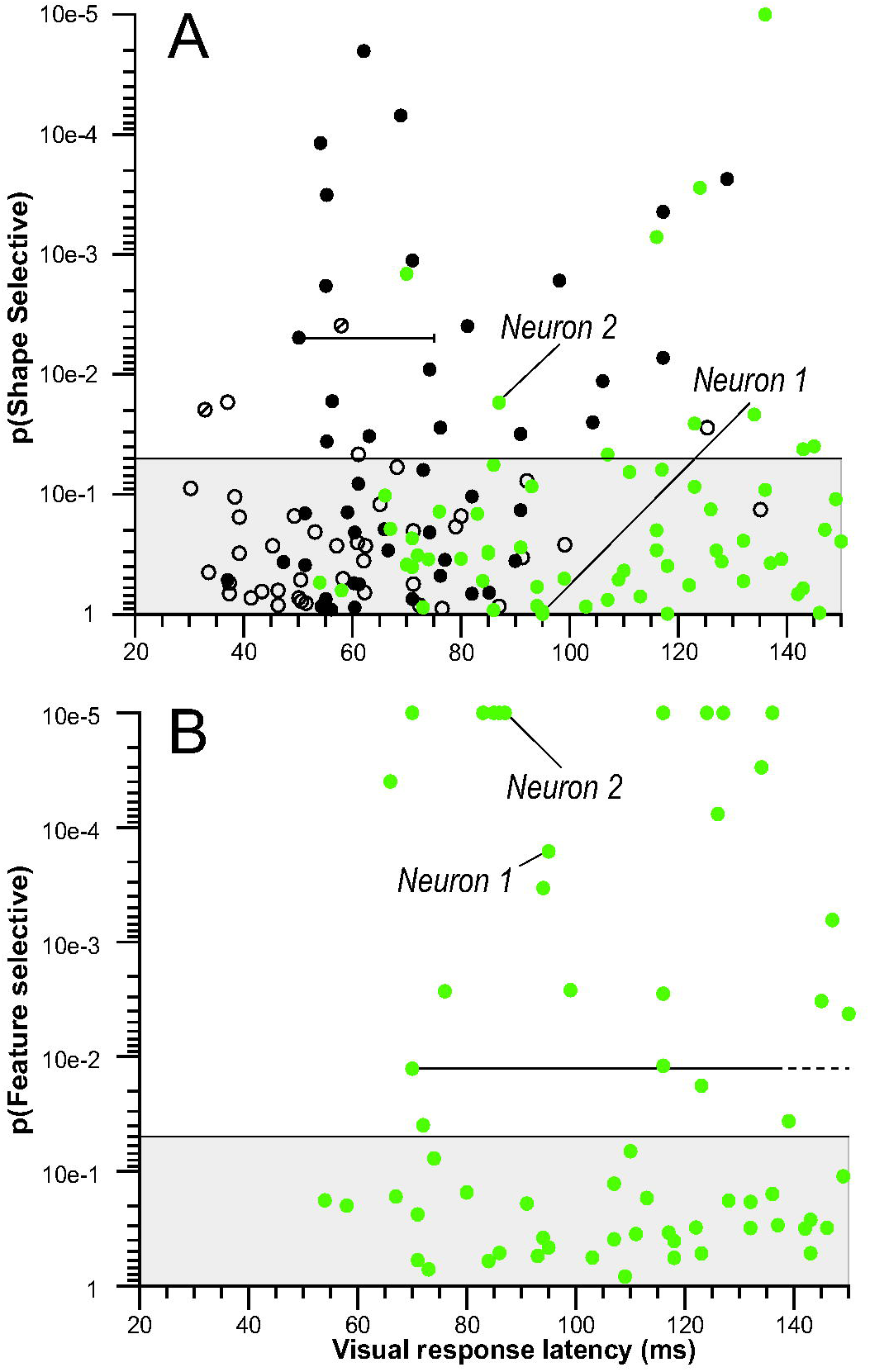
Relationship between feature selectivity and visual latency compared for neurons sampled in this study (green) and those reported previously in control monkeys that performed search with variable color assignments (open black circles) and experimental monkeys that performed search with constant color assignments (filled black circles) (Bichot et al. 1996). The probability of the response to the singleton in the receptive field being the same as the response to a distractor in the receptive field during the first 25 ms (A) and 100 ms (B) is plotted as a function of visual response latency. Horizontal lines indicate analysis window. In (B) the dashed portions of the line indicate that the 100 ms analysis window extends beyond the range of the plot. The shaded region indicates nonsignificant probability values greater than 0.05. In the previous study, of the 43 neurons from control monkeys, 39 fell in the nonsignificant area, two responded preferentially to the target, and two responded preferentially to the distractors of the search array field (marked by diagonal lines). In contrast, 21 of 47 neurons recorded from the experimental monkeys exhibited significantly greater initial responses when the singleton fell in the receptive field, and none showed the opposite effect. In the current study, of 124 neurons sampled, 16 showed shape selectivity in the first 25 ms and 30 in the first 100 ms. Example neurons 1 and 2 are identified as N1 and N2.

### Relation of Feature Selection to Spatial Selection

The serendipitous discovery of orientation sensitivity in FEF offered an opportunity to relate these observations to previous findings (Thompson et al., 1996; Murthy et al., 2001; Sato & Schall, 2003; Schall 2004). We performed the following sequence of analyses. To report the findings most clearly and concisely, we introduce a nomenclature to distinguish the categories of neurons, the types of trials and the timing measures. First, as previously, we distinguish singleton selection time (SST) from saccade endpoint selection time (EST). Second, we distinguish whether measures were obtained in correct or error trials with left subscript, e.g., _Corr_EST and _Err_EST. Third, we distinguish whether measures were obtained in pro- or anti-saccade trials with right subscript, e.g., _Corr_EST_Pro_ and _Corr_EST_Anti_. Finally, we distinguish whether the measure was obtained in trials with congruent, incongruent, or neutral search arrays with right superscript, e.g., _Corr_EST^C,I^_Pro_ and _Corr_EST^C,I^_Anti_. The absence of a particular superscript or subscript implies that the measure was obtained over all possible groups. The authors appreciate the complexity of this nomenclature, which is in keeping with that of more mature scientific fields such as chemistry, molecular biology, and physics that require non-intuitive but detailed nomenclatures and symbols.

In the first analysis, responses during pro- and anti-saccade trials were assessed for the feature selective and the non-feature selective neurons to identify SST and EST as measured previously (Sato & Schall, 2003) (Fig. 5A). In pro-saccade trials, the average response became greater when the singleton was in the RF relative to when it was opposite the RF, replicating Sato & Schall (2003) and numerous other studies describing target selection in FEF during search (e.g., Bichot et al., 2015; Buschman & Miller, 2007; Glaser et al., 2016; Keller et al., 2008; McPeek, 2006; Mirpour et al., 2019; Monosov et al., 2010; Monosov & Thompson, 2009; Phillips & Segraves, 2009; Pouget et al., 2009; Scerra et al., 2019; Schall et al., 1995; Schall & Hanes, 1993; Thompson et al., 1996; Wardak et al., 2006; Zhou & Desimone, 2011). Conversely, in anti-saccade trials, the average response across the sample of feature selective neurons became greater when the endpoint of the saccade was in the RF relative to when the singleton was in the RF. Similar results were found for the non-feature-selective neurons (Fig. 5B).

**Figure 5.**
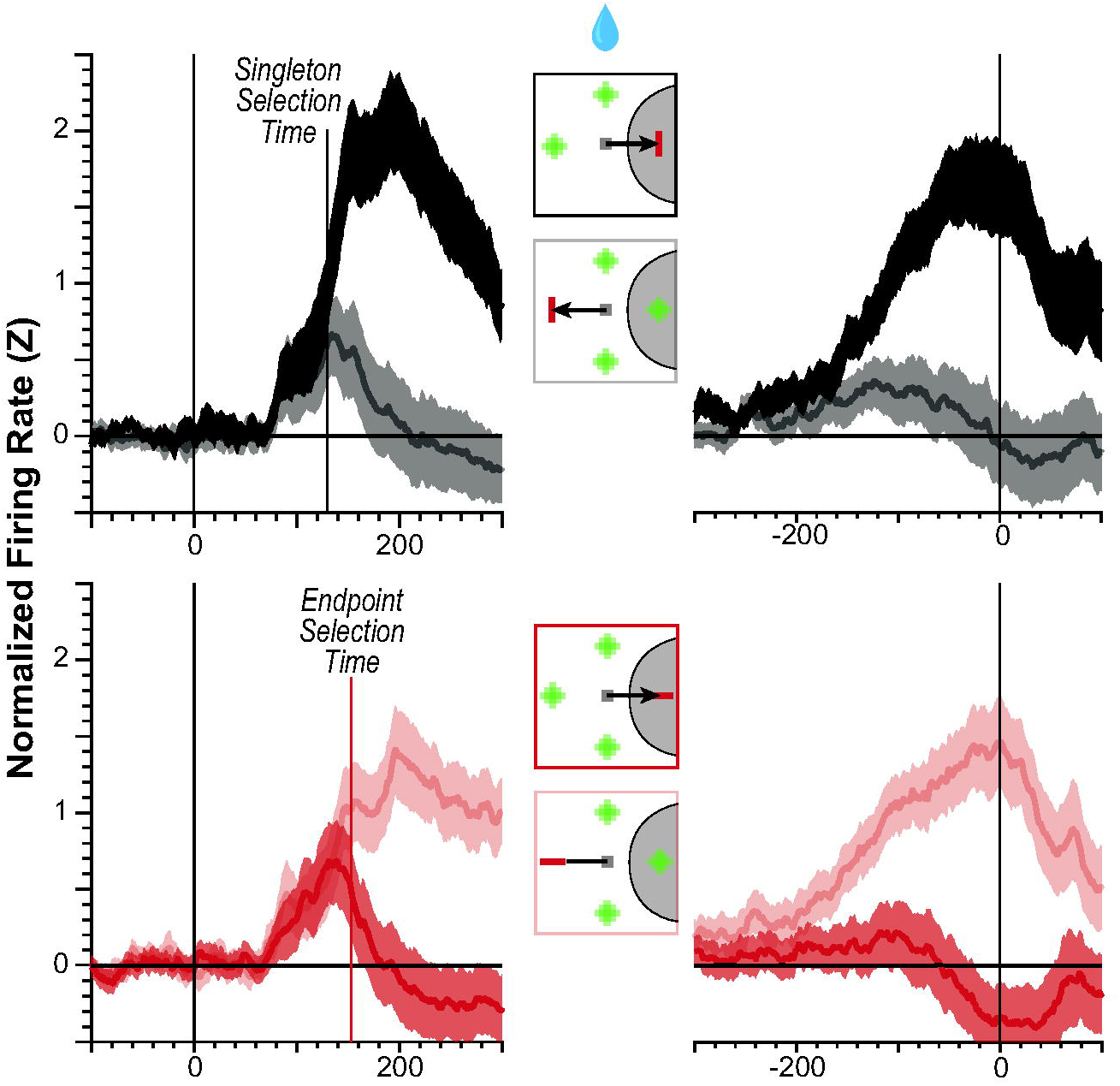
Singleton and saccade endpoint selection. (A) For the 30 feature selective neurons, average normalized SDF when the singleton appeared in (dark) or opposite (light) the RF during interleaved pro- (top) and anti-saccade (bottom) trials aligned on array presentation (left) and on saccade initiation (right). Insets illustrate the locations and orientations of the singleton and possible horizontal, square, or vertical distractors relative to RF (gray arc) plus the reward earned (drop icon) for each SDF. SST measures when the SDF for the singleton in the RF exceeds the SDF for a distractor in the RF. EST measures when the SDF for the anti-saccade endpoint opposite the RF exceeds the SDF for the singleton in the RF.

These results generally replicate previous observations (Sato & Schall, 2003); however, the absence of SST during anti-saccade trials was unexpected. The monkey’s performance strategy resulted in low accuracy for Anti^N^ and Anti^I^ trials. Hence, the absence of SST_Anti_ is consistent with a failure to focus attention on the singleton appropriately. Further, the aspect ratio of the stimuli used in this study was greater than that used by Sato & Schall and so was more easily discriminable from central fixation. However, when RTs were longer, due either to more deliberate focusing of attention on the singleton or overall slowing of processing, SST preceded EST during anti-saccade trials (Fig. 6). Therefore, the overall pattern of neural modulation observed in FEF is consistent with the performance data indicating that the monkey divides attention among vertical items, sacrificing accuracy for speed.

**Figure 6.**
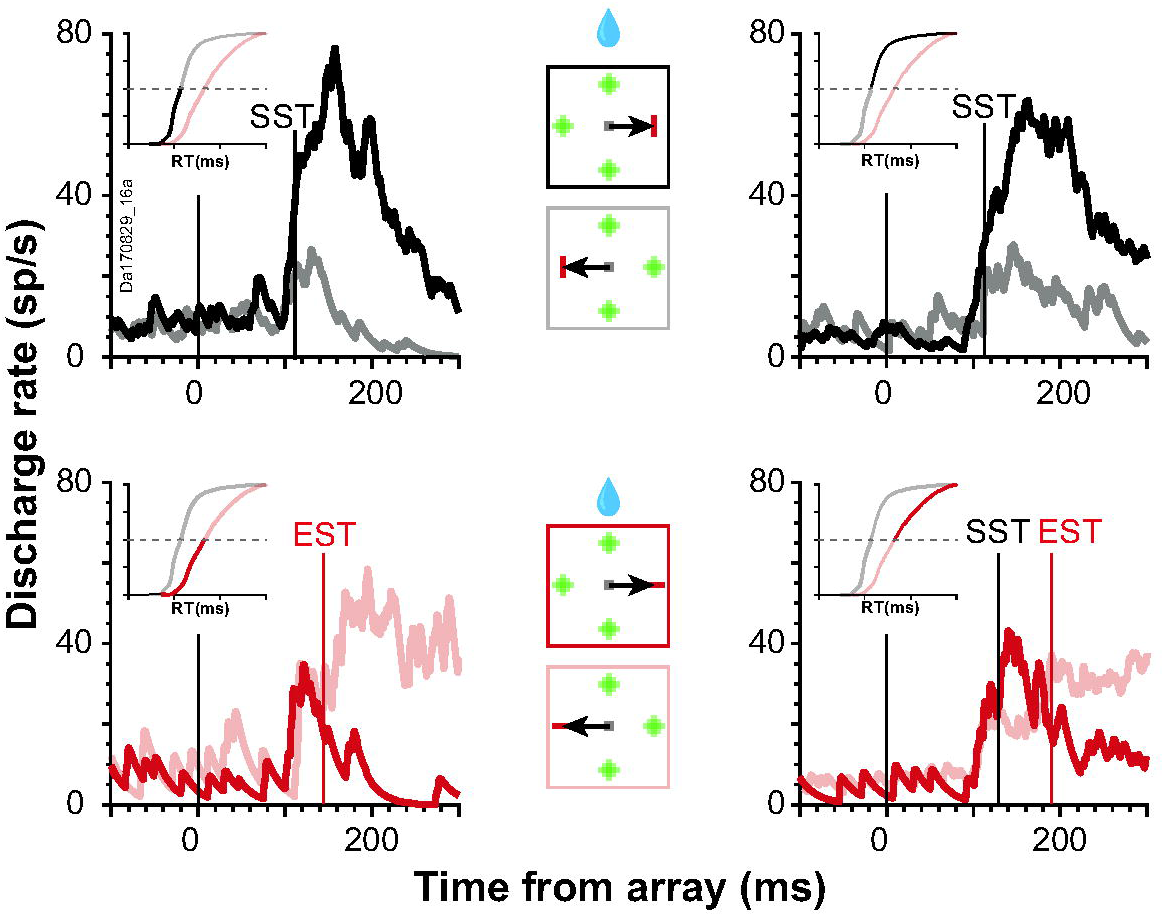
Singleton and saccade endpoint selection across response time. Representative neuron illustrating variation of SST and EST for shortest (left) and longest (right) RT (highlighted in inset cumulative RT distributions). In pro-saccade trials, SST does not vary with RT. In anti-saccade trials, SST was manifest in long but not short RT trials, followed by EST. Conventions as in Figure 4.

Across the sample of feature selective neurons, SST measured in pro-saccade trials (_Corr_SST_Pro_) preceded EST measured in anti-saccade trials (_Corr_EST_Anti_) Average values for these and all subsequent temporal indices ± SEM are found in Table 1. Statistical tests on all pairs of distributions are found in Table 2.

**Table 2.**
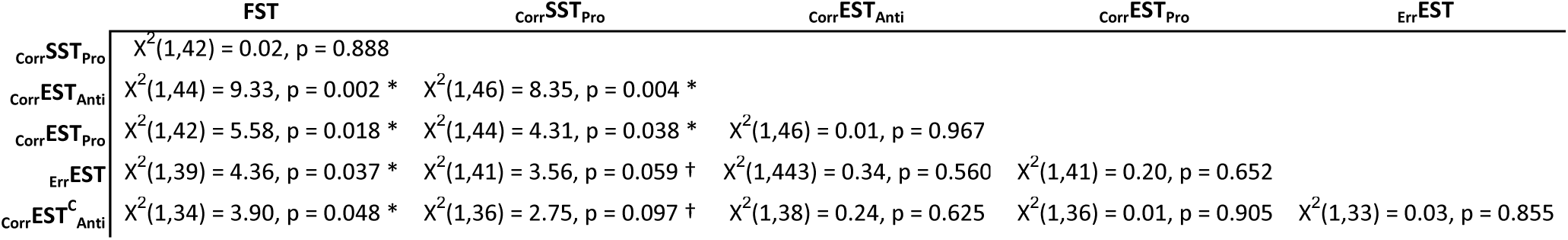
Selection time comparisons. The distribution of each selection time was compared to the distribution of each other selection time using a Kruskal-Wallis test. The Χ^2^ value, degrees of freedom, and *p* value of each pairwise test is shown. Because the tests are symmetric, only the lower diagonal is shown. Values that trend toward significance (*p* < 0.10) are marked with a dagger (†). Values that reach significance (*p* < .05) are marked with an asterisk (*).

Having established that these relationships replicate previous observations (Sato & Schall, 2003), we can now explore the relationship of the new measure FST to SST and EST measured in the different types of trials. FST was not significantly different than _Corr_SST_Pro_. In contrast, FST was significantly earlier than _Corr_EST_Anti_.

The simultaneity of FST with _Corr_SST_Pro_ entails that they index a common process. If so, then FST can inherit the interpretation of SST. Accordingly, we conjecture that FST indexes the process of stimulus selection through attention allocation and not saccade endpoint selection.

The second analysis assessed how feature selection was related to spatial selection of locations other than the singleton or saccade endpoint. This was accomplished by contrasting responses of feature-selective neurons to fixated and non-fixated stimuli. Fig. 7A compares the activity of the two example neurons and of the sample of feature-selective neurons to vertical distractors in the RF that were not fixated, activity preceding correct pro-saccades to the vertical singleton in the RF, and activity when unchosen square or horizontal distractors were in the RF. Responses were greater when the vertical color singleton in the RF attracted a gaze shift relative to when a vertical distractor in the RF was not fixated, replicating the well-known enhancement effect (Goldberg & Bushnell, 1981). By comparing discharge rates when an unfixated, irrelevant vertical distractor was in the RF and when the fixated vertical color singleton was in the RF, we measured *endpoint selection time* for pro-saccades (_Corr_EST_Pro_).

**Figure 7.**
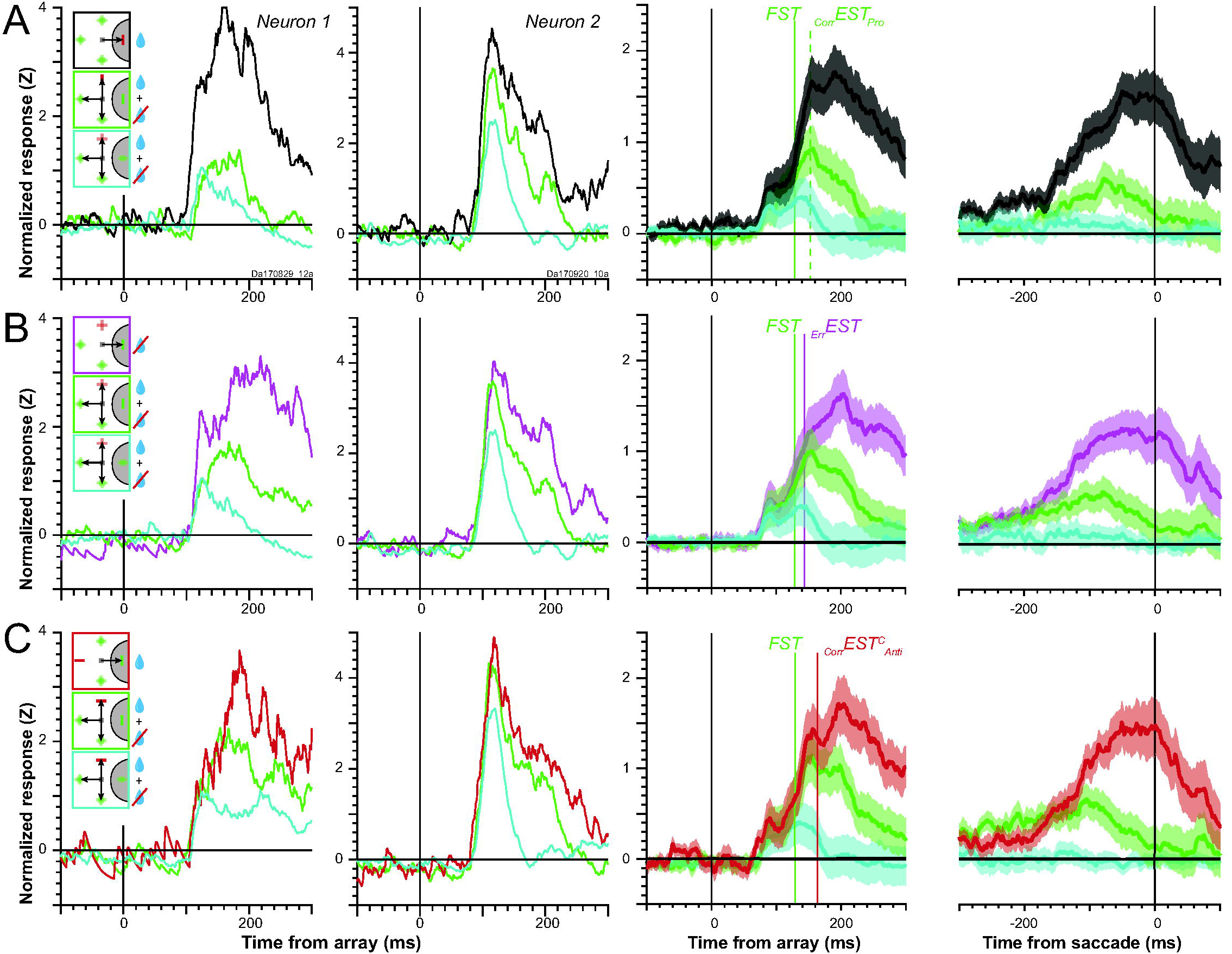
Distinction of feature selectivity from saccade selection. Normalized firing rates for neuron 1 (1^st^ column) and neuron 2 (2^nd^ column) aligned on array presentation, plus mean normalized SDF ± SEM of feature selective neurons aligned on array presentation (3^rd^ column) and on saccade initiation (4^th^ column). (A) Activity associated with irrelevant vertical (green), non-vertical (cyan), and the singleton in the RF (black) demonstrate enhancement associated with correct saccade selection, which distinguishes FST from _Corr_EST_Pro_. (B) Activity on pro- and anti-saccade trials associated with irrelevant vertical (green), non-vertical (cyan), and incorrectly selected vertical distractors in the RF (magenta) demonstrate enhancement associated with errant saccade selection, which distinguishes FST from _Err_EST. (C) Activity on anti-saccade trials associated with irrelevant vertical (green), non-vertical (cyan), and correctly selected vertical distractor in the RF (red) demonstrate enhancement associated with anti-saccade selection, which distinguishes FST from _Corr_EST^C^_Anti_.

The time _Corr_EST_Pro_ identifies when the endpoint of the upcoming pro-saccade is specified by feature-selective neurons. This is a new measure. It is distinct from EST defined by Sato and Schall (2003), or _Corr_EST_Anti_ described above because it was not calculated from anti-saccade trials. Across the sample of feature selective neurons, _Corr_EST_Pro_ was significantly later than FST and _Corr_SST_Pro_, but was not different from _Corr_EST_Anti_.

The third analysis tested whether _Corr_EST_Pro_ was due only to the difference in color between the fixated and unfixated vertical items. This was accomplished by contrasting responses when an incorrect saccade was made to a vertical distractor in the RF relative to the unfixated vertical distractor (Fig. 7B). The response to the fixated vertical distractor was greater than the response to the un-fixated vertical distractor. This replicates multiple previous findings that saccade endpoint errors during visual search arise when FEF neurons treated a distractor as if it were the target (Heitz et al., 2010; Reppert et al., 2018; Thompson et al., 2005). We identify the time when this occurs as *endpoint selection time* for errors (_Err_EST). Across the sample of feature selective neurons, _Err_EST was significantly later than FST and trended toward being later than _Corr_SST_Pro_, but was not different than _Corr_EST_Anti_ or _Corr_EST_Pro_.

The fourth analysis tested whether the responses of feature-selective neurons varied across trial context. This was accomplished by comparing the responses observed with correct anti-saccades to the vertical item and responses to irrelevant vertical and non-vertical distractors (Fig. 7C). This analysis compared only items of the same color. Both example neurons produced most activity associated with fixated vertical stimuli in the RF relative to un-fixated vertical distractors, and least activity with square or horizontal distractors in the RF. Across the sample of feature selective neurons, the *endpoint selection time for congruent anti-trials (*_Corr_EST^C^_Anti_) was significantly later than FST but was not different than_r_SST_Pro_, _Corr_EST_Anti_, _Corr_EST_Pro_, or _Err_EST.

These analyses assess the temporal aspects of attention allocation and endpoint selection. Fig. 7 shows three conditions in which vertical items were fixated: correct Pro trials, incorrect saccades to vertical items, and correct Anti^C^ trials. These were used to identify _Corr_EST_Pro, Err_EST, and _Corr_EST^C^_Anti_, respectively. In a fifth analyses, the magnitude of response in three conditions were compared at three time windows: 100 to 150 ms after array onset (around the time of FST and _Corr_SST_Pro_), 150 to 200 ms after array onset (around the time of EST), and - 25 to 25 ms from saccade initiation (Fig. 8). The magnitude of the responses did not differ in the early visual time window (F(2,87) = 0.022, p = 0.9774), the late visual time window (F(2,87) = 0.077, p = 0.9263), or around the saccade (F(2,87) = 0.106, p = 0.8994). In short, responses were identical if a saccade was made toward a vertical item in the RF, regardless of context or whether such a saccade was correct or incorrect.

**Figure 8.**
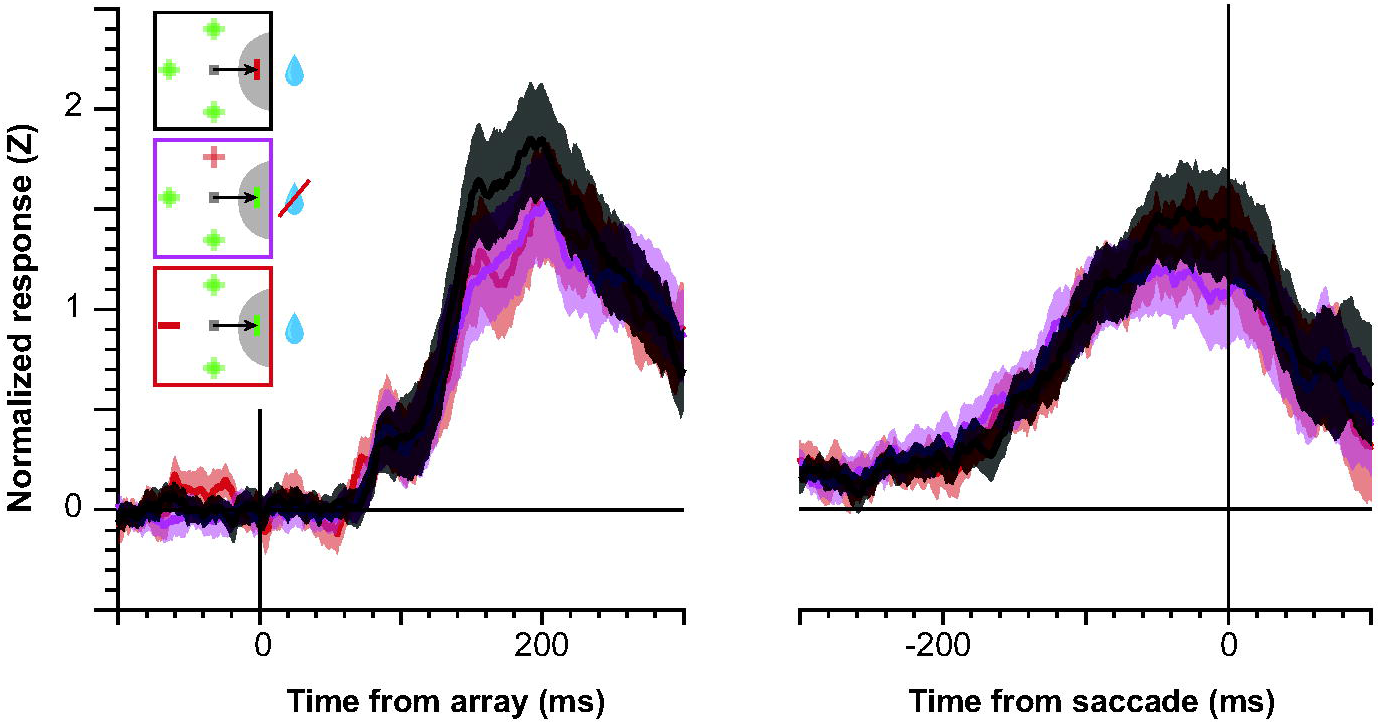
Magnitude of response during saccade selection. Mean normalized SDF ± SEM of feature selective neurons aligned on array presentation (left) and on saccade initiation (right). Activity associated with correct Pro saccades into the RF (black), incorrectly selected vertical distractors in the RF (magenta), and correct Anti^C^ saccades into the RF (red) do not differ, showing that this population does not differentiate type of saccade if a saccade is to be made.

### Variation of Modulation Times in Relation to RT

Previous research using this task distinguished neurons by measuring whether SST and EST were synchronized on array presentation or varied with RT (Sato & Schall, 2003; Schall 2004). We performed the same analysis for these data, calculating FST, _Corr_SST_Pro_, _Corr_EST_Anti_, _Corr_EST_Pro_, _Err_EST, and _Corr_EST^C^_Anti_ in the fastest and slowest 50% of trials. The difference in selection times divided by the interquartile range of the RTs could range between 0.0 (synchronized on array presentation) to 1.0 (synchronized on saccade initiation).

The proportion of RT accounted for by variation in selection times are shown in Fig. 9. We found that this proportion was not different than 0.0 for FST (t(13) = −0.49, p = 0.683) or _Corr_SST_Pro_ (t(18) = 0.91, p = 0.377). In terms of Bayes Factors (Rouder et al., 2009) we found moderate evidence that *FST* (BF = 0.22) and _Corr_SST_Pro_ (BF = 0.28) account for no variability of RT. In other words, the state indexed by FST and _Corr_SST_Pro_ arises at a time synchronized on array presentation.

**Figure 9.**
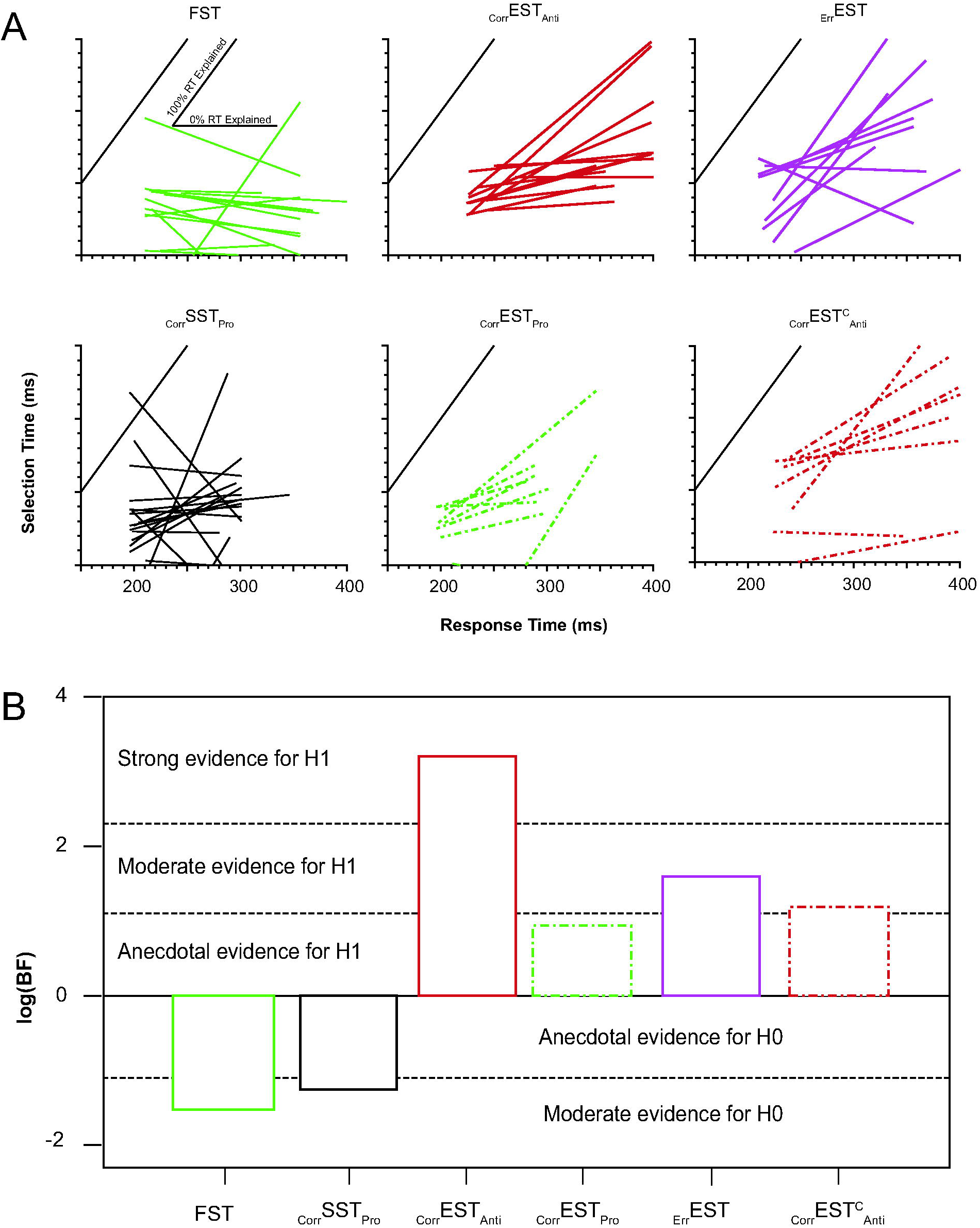
Chronometry of feature selection, singleton selection, and endpoint selection in relation to response time. (A) Selection times for faster and slower RT groups plotted as a function of the mean RT of each group. Each line corresponds to one neuron with a measurable selection time in both RT groups. The slope indicates the contribution of each selection time to RT. Inset in top left subplot (FST) illustrates range of possible influences of selection times on RTs. Selection times could be synchronized on array presentation and invariant with respect to RT (0% RT explained) or synchronized on saccade presentation (100% RT explained). Colors as in Fig. 6. Dashed lines indicate measures from non-feature-selective cells. (B) Bayes factors from statistical test of the slopes of each selection time relative to RT. Bayes factors less than 1 (log values less than 0) indicate evidence for the null hypothesis (H0) that the distribution mean is equal to 0. Bayes factors greater than 1 (logs greater than 0) indicate evidence for the alternate hypothesis (H1) that the distribution is greater than 0. Levels of evidence defined by the Bayes factor are indicated. Line and color assignments as in Fig 6. We found moderate evidence supporting the hypothesis that FST and _Corr_SST_Pro_ are synchronized on array presentation and not on saccade initiation. On the other hand, we found strong evidence that _Corr_EST_Anti_, anecdotal evidence that _Corr_EST_Pro_, and moderate evidence that _Err_EST and _Corr_EST^C^_Anti_ were not synchronized on array presentation nor saccade initiation.

In contrast, variation in all measures of endpoint selection in feature-selective cells accounted for a significant fraction of variation of RT. With strong evidence rejecting the null hypothesis (BF = 24.62), a significant proportion of the variation of RT was accounted for by variation in _Corr_EST_Anti_ (t(13) = 3.92, p = 0.002). At a moderate level of evidence, a significant proportion of the variation of RT was accounted for by variation in _Err_EST (t(9) = 3.22, p = 0.011, BF = 5.64) and _Corr_EST^C^_Anti_ (t(7) = 2.95, p = 0.021, BF = 3.29). At an anecdotal level of evidence, a significant proportion of the variation of RT also was accounted for by variation of _Corr_EST_Pro_ (t(8) = 2.71, p = 0.027, BF = 2.55).

Although the measures of EST account for some RT variability, the average proportion of RT explained across all significant relationships is 24.8%. The additional RT variability will be accounted for by response preparation processes subsequent to EST and not included in these data.

### Neural Chronometry of Feature and Spatial Selection

The various distinct response modulations reveal a temporal sequence of operations in FEF accomplishing this visual search task (Fig. 10; Table 2). Following array presentation, the first state transition is indexed by the response of visually responsive neurons after a characteristic latency. The next state transition was indexed by FST, which coincided with _Corr_SST_Pro_. The state indexed by _Corr_SST_Pro_ has been identified with the allocation of visual attention on the singleton based on its salient visual attribute to encode the stimulus-response rule (Sato & Schall, 2003; Schall 2004). The discovery of feature-selection arising concomitantly with _Corr_SST_Pro_ reported here suggests that the monkey divided visual attention among the vertical items in the array. The allocation of spatial visual attention to spatially separated, noncontiguous items in a search array has been demonstrated (e.g., Bichot et al., 1999; Dubois et al., 2009). The next state transition was indexed by EST. The state indexed by EST has been identified with the specification of the endpoint of the saccade. Being different in time and relationship with RT, it is a state different from that identified by _Corr_SST_Pro_ (Sato & Schall, 2003; Schall 2004) and likewise distinct from the presaccadic build-up of movement related neurons (Woodman et al., 2008), which accounts for the remainder of the variation of RT.

**Figure 10.**
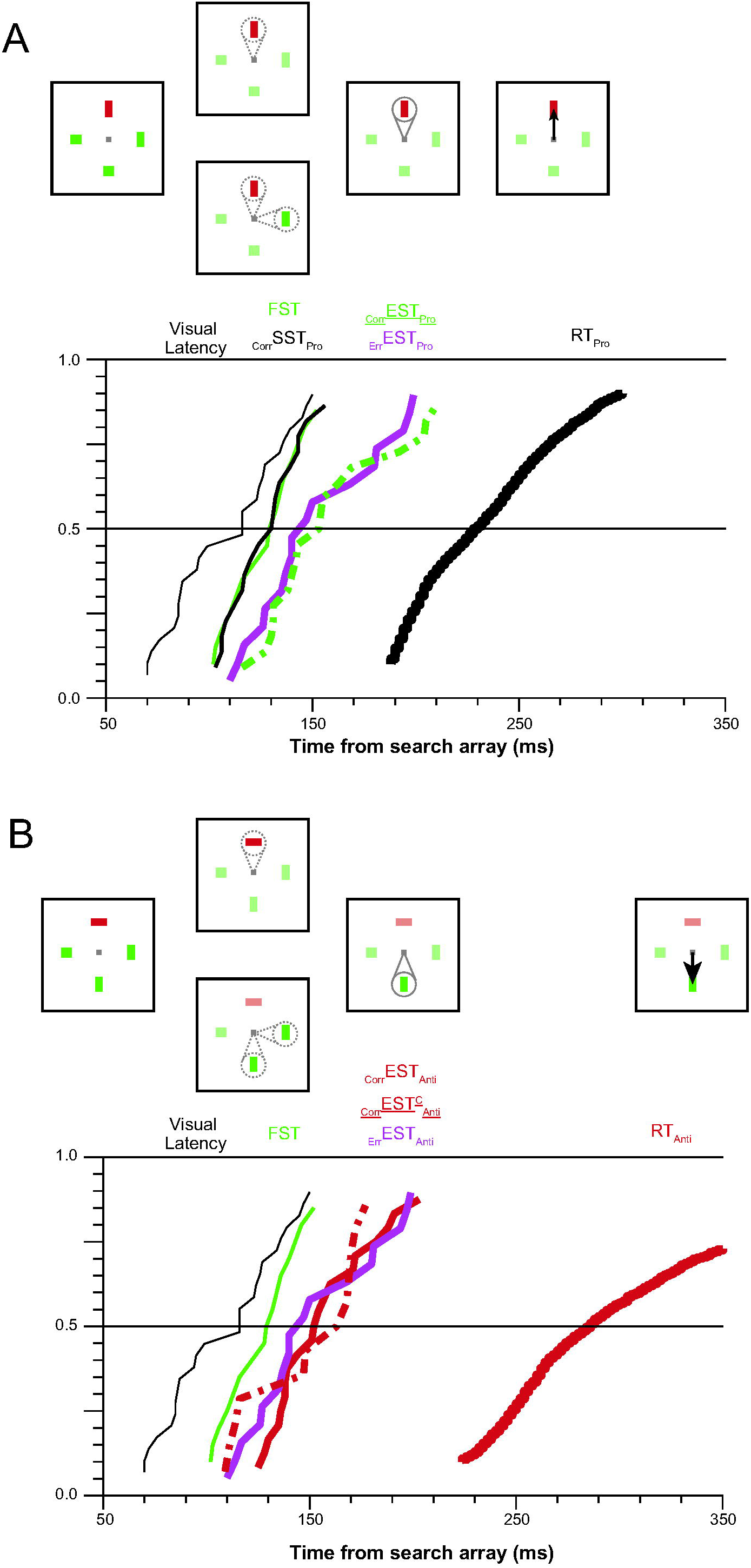
Distributions of feature selective processes. (A) Diagrams showing sequence of states during pro-saccades (top). The hypothesized spotlight of attention is shown in gray lines and a saccade is indicated by a solid arrow. Cumulative distributions of selection time metrics alongside visual response latency and Pro RT distribution (bottom). The colors are the same as the diagrams and previous figures and labeled above the plot boundary. Line thickness increases as stages become further from array onset and closer to RT. (B) Diagrams showing sequence of states during anti-saccades (top) and cumulative distributions of selection time metrics (bottom).

## DISCUSSION

The present study demonstrates two primary findings: (1) besides color (Bichot et al., 1996), shape selectivity can arise in FEF when strategies commit feature attention and (2) this feature selectivity, which seems associated with divided attention, is functionally distinct from the selection of the saccade endpoint. The first finding may seem at odds with the perspective that FEF selects targets regardless of the feature that identifies a stimulus as that target. However, adaptive performance strategies can explain this anomaly. Strategies are revealed by analyzing the responses made on error trials and RT in all trials. The increased prevalence of error saccades to vertical stimuli and the fastest RT to vertical stimuli reveals a priority for locating vertical stimuli.

The results are based on data obtained from a single monkey. Nevertheless, we believe they are reliable and interpretable for the following reasons. First, the observation of feature selectivity in FEF replicates previous findings (Bichot et al. 1996; Peng et al. 2008). A similar predisposition for motion direction has been described in the superior colliculus of monkeys performing a motion discrimination task with fixed stimulus-response mapping (Horwitz et al., 2004). The unexpected but clear robustness of this phenomenon should engender confidence in the replicability of the current observations. Second, the distinction of singleton selection and endpoint selection replicates previous findings (Sato & Schall, 2003; Schall 2004). Such replication should increase confidence in the interpretability of the new findings. Finally, the novel observation of a distinct endpoint selection in pro-saccade trials is statistically robust, conceptually novel, and theoretically important. While we are confident that another monkey could be trained into this state, we judge that effort is better invested in more novel research goals. Indeed, we have discovered that the second monkey, trained without the opportunity to experience the confounds, employs a qualitatively different strategy to perform this task (Lowe et al., 2019).

### Possible Sources of Feature Selection in FEF

We do not know whether the shape selectivity we observed is intrinsic to FEF, imparted by other prefrontal areas, inherited from earlier visual areas, or manifest from broad associations of stimulus, action, and reward. We consider each hypothesis below.

The hypothesis that feature selectivity is intrinsic to FEF runs counter to the framework of FEF as an area that contains a salience or priority map regardless of features defining salience or priority (Thompson & Bichot, 2005). However, some studies have reported differential activity to stimuli defined by features whose identities do not dictate different stimulus-response rules (Ferraina et al., 2000; Peng et al., 2008; Xiao et al., 2006). Mohler et al. 1973 reported 6% of FEF neurons (12.5% of those with visual responses) responding differently according to direction of motion or color. Peng and colleagues (2008) found that even during a passive fixation task a quarter of FEF neurons had responses that differed according to the form of the presented stimuli. These differences occurred at most 12 ms after the initial visual transient. This short delay between visual response onset and feature selectivity is consistent with the selectivity for color found previously (Bichot et al. 1996). However, the shape selectivity presented here was not as immediate. This may be due to the nature of the tasks across studies in that there are unbalanced reward contingencies of nonpreferred stimuli in the present study whereas all stimuli were evenly rewarded in the passive fixation and delayed match to sample tasks used by Peng et al. It is notable that the proportions of feature selective neurons found by Peng et al. are similar to those found in the present data, but are fewer than those found by Bichot et al. (1996). This could be due to differences in complexity of the stimuli, nature of the task, or sampling of units.

The hypothesis that feature selectivity in FEF can be imparted by another prefrontal area is motivated by the recent description of a ventral prearcuate area (Bichot et al. 2015), which has dense connections with FEF (Huerta et al., 1987). Neurons in this area have differential responses to complex visual stimuli during detection and delayed search tasks, and this feature selectivity preceded the selection of a saccade endpoint (Bichot et al., 2015). However, direct comparison between this and the current study is challenged by differences in experimental design and particular observations. For example, their target item was cued before array presentation and so was held in working memory, but our target item in this study was a long-term memory trace. Also, neurons in the ventral prearcuate area exhibited feature selectivity at approximately the same time as FEF, and the spatial selectivity identified in FEF was earlier than that observed in the present data (_Corr_SST_Pro_). Further research is needed, therefore, to clarify whether FEF receives feature information primarily from this area, or both areas have common inputs and process feature information in parallel.

The hypothesis that feature selectivity in FEF is inherited from feature selective responses earlier in the visual stream is motivated by the connections between FEF and effectively all extrastriate visual areas (Schall et al. 1995; Markov et al. 2014). V4 is one likely source because the neurons are selective for color (Schein & Desimone, 1990; Zeki, 1980; Zeki, 1973) and shape (Desimone & Schein, 1987; Pasupathy & Connor, 1999). In the previous (Bichot et al. 1996) and current study, neither color nor shape selectivity were carried by the FEF neurons with the shortest visual latencies. This is consistent with color and shape information arriving in relatively longer latency afferents (e.g., Schmolesky et al., 1998). Evidence from simultaneous recordings in FEF and V4 demonstrate an association of visual neurons in FEF with V4 (Gregoriou et al., 2012) and feature selectivity in V4 preceding FEF selective modulation (Zhou & Desimone, 2011). Further research is needed, though, to understand the interplay of feature selectivity and attentional modulation between FEF and extrastriate visual areas (Zhou et al., 2011; see also Monosov et al., 2010).

The hypothesis that feature selectivity in FEF is manifestation of the association of strategy and reward is motivated by well-known reports that visual responses in FEF are modulated by reward expectation (Glaser et al., 2016) or magnitude (Ding & Hikosaka, 2006). Parallel modulation is observed broadly in the visuo-motor network (e.g., Griggs et al., 2018; Platt & Glimcher, 1999; Sugrue et al., Newsome, 2004; Yamamoto et al., 2013). In human studies, both reward probability and magnitude have been shown to influence behavior. Della Libera & Chelazzi, (2009) found that by associating meaningless shape stimuli with high, low, or neutral reward in a practice phase resulted in facilitation or interference of response times, depending on task conditions. Similarly, attentional biases emerge when color stimuli are associated with high or low reward, whether or not participants are aware of the stimulus-reward associations (Kiss et al., 2009; Kristjánsson et al., 2010). These associations do not require physical salience as they are present with stimulus configurations that have only reward histories to differentiate stimuli and for which rewarded features are not the basis for selection (Anderson et al., 2011). These findings suggest that stimulus-reward associations can be learned and combined with physical salience to form an integrated priority map (Awh et al., 2012). These reward associations manifest themselves in neural activity (Anderson, 2016). The tail of the caudate is sensitive to learned reward associations (Anderson et al., 2014). Learned value associations are reflected in BOLD signaling in attentional visual areas such as parietal cortex (Anderson et al., 2014) and are reflected in shifts of ERPs indexing attentional selection such as the N2pc (Kiss et al., 2009).

In conjunction search FEF neurons respond maximally when the correct saccade target is in the RF (Bichot et al., 2001; Ogawa & Komatsu, 2006) but also show larger responses to a distractor that shares a feature with the correct saccade target than a distractor that shares no features (Bichot et al., 2001). Similarly, FEF neurons respond more when a distractor that was the target on the previous session is in the RF than a distractor that shares no features with the current saccade target. This demonstrates that FEF neurons can differentially respond to features that are remembered to be rewarded even when not presently rewarded. Reward associations, specifically the lack thereof, can also participate in distractor suppression (Cosman et al., 2018). In a search task with salient distractors that “capture” attention (Theeuwes, 1991) two monkeys overcame capture with training and produced equal performance when the color singleton distractor was present or absent. Neurons recorded from those two monkeys showed a reduction in firing rate when the salient distractor was in the RF compared to a non-salient distractor was in the RF. Because the salient distractors were never a saccade target, but were nevertheless distinguishable from the other distractors, responses to them can be more actively and immediately suppressed than the other distractors. Bichot and colleagues (2001) also tested neural responses during a search task with a salient distractor and did not find distractor suppression. However, the monkeys in that study were behaviorally affected by the singleton distractor and thus distractor suppression may not be expected. Further, the neurons analyzed by Bichot and colleagues were movement neurons whereas those analyzed here and by Cosman et al. had visual responses. This difference in neuron type may also explain the differences in results.

Interestingly, the third monkey in the study by Cosman and colleagues that was unable to overcome attentional capture was the same monkey Da whose data are reported here. Neurons from this monkey did not show such distractor suppression. Notably, this monkey also had neurons that retained an initial nonspecific visual response whereas monkeys A and C did not have such a response during the color singleton search task. Such an initial visual response is reduced in FEF neurons when stimuli are not saccade targets (or, alternatively, enhanced when they are saccade targets) in both search tasks (Thompson et al., 1997) and in single stimulus presentations (Goldberg & Bushnell, 1981; Mohler & Wurtz, 1976; Schall et al., 1995). In the case of monkeys A and C, the stimuli whose colors were not the target color were never correct saccade targets and can thus be discounted and would have attenuated nonspecific responses to these stimuli, and this attenuation could be complete such that there is no such response. In the case of Da, square and horizontal stimuli were correct saccade endpoints on a subset of anti-saccade trials, thus they are still associated with reward to some degree and thus may require the retaining of the nonspecific visual transient.

### Processing Operations and Neural Chronometry

We replicated the previous finding of distinct operations mediated by visually responsive neurons selecting a conspicuous stimulus and selecting the endpoint of the saccade (Sato & Schall, 2003). The prior experiment did this by contrasting modulation in pro- and anti-saccade trials. The current experiment did this, innovatively, by contrasting modulation to preferred and non-preferred features and to fixated and non-fixated items among identified neurons exhibiting feature selectivity even for stimuli that should not be and were not selected. Specifically, we demonstrated quantitative differences between two measures of neural modulation: stimulus selection, indexed by FST and _Corr_SST_Pro_, and saccade endpoint selection, indexed by EST. The chronometric distinction between singleton selection and endpoint selection in both pro- and anti-saccade trials and the simultaneity of EST on pro- and anti-saccade trials having very different RT validates the conceptual distinction between these operations. These neural measures index some of the computational operations occupying response time in this task (Donders, 1969).

The delay between EST and saccade initiation identifies another operation preceding saccade initiation. This operation has been identified psychologically as response preparation and neurally as the presaccadic build-up of movement related neural activity, which does not occur until information about target items becomes available (Woodman et al., 2008) and is identified with the accumulation of sensory evidence (Purcell et al., 2010, 2012; Servant et al., 2019). The final saccade initiation operation is accomplished by competitive interactions between movement cells (Purcell et al., 2010, 2012). The time required for this competition resolution explains the additional time necessary for anti-saccades compared to pro-saccades. The relationship between stimulus selection, endpoint selection, and saccade preparation has been investigated in monkeys (Juan et al., 2004; Katnani & Gandhi, 2013) and humans (Juan et al., 2008).

To verify the existence and elucidate the properties of these distinct operations and stages, and to resolve different explanations for causal manipulations, further research should employ the powerful logic of selective influence in factorial experimental designs (Sternberg, 2001; Townsend & Nozawa, 1995) with joint measures of mental and neural chronometry.

## Acknowledgements

The authors thank J. Elsey, M. Fuertado, M. Maddox, S. Motorny, J. Parker, M. Schall, C.R. Subraveti, and L. Toy for animal care and other technical assistance, S. Errington, T. Reppert, A. Sajad, and J. Westerberg for helpful discussions regarding the work.

